# The effect of heavy metal contamination on human and animal health in the vicinity of a zinc smelter

**DOI:** 10.1101/459644

**Authors:** Xiaoyun Shen, Yongkuan Chi, Kangning Xiong

**Author notes:** Corresponding author. E-mail addresses. These authors contributed equally to this work (Xiaoyun Shen, Yongkuan Chi).

## Abstract

A diagnosis of heavy metal poisoning in sheep living on pasture in the vicinity of a zinc smelter was made based on laboratory tests and clinical signs in livestock in the Wumeng mountain area of China. Heavy metal contamination has generated serious harm to the health of local farmers after passing through the food chain. The levels of copper, zinc, cadmium, and lead in irrigation water, soil, forages, and animal tissues were measured in samples taken from within the vicinity of a zinc smelter and control samples. Heavy metal concentrations in foods (corn, rice, and wheat) and human tissues (blood and hair) from local farmers living in affected areas and controls were also determined. Hematological values were determined in human and animal samples. The copper, zinc, cadmium, and lead concentrations in irrigation water, soils, and forages were markedly higher than the levels in healthy pastures. Cadmium and lead concentrations were 177.82 and 16.61 times greater in forages than controls, respectively, and 68.71 and 15.66 times greater in soils than controls, respectively. Heavy metal concentrations in food (corn, rice, and wheat) in affected areas were markedly higher than in the control samples. Cadmium and lead concentrations in the tissues of affected sheep were markedly higher than in control animals (*P< 0.01*). Cadmium and lead concentrations in blood and hair samples from affected farmers were markedly higher than the control samples (*P < 0.01*). The occurrence of anemia in affected persons and animals followed a hypochromic and microcytic pattern. The intake of cadmium and lead was estimated according to herbage ingestion rates. It was found that the levels of cadmium and lead accumulated in sheep through the ingestion of vegetation growing in the sites closest to the zinc smelter were approximately 3.36 mg Cd/kg body wt./day and 38.47 mg Pb/kg body wt./day. This surpassed the fatal dosages for sheep of 1.13 mg Cd/kg body wt/day and 4.42 mg Pb/kg body wt./day. Serum total antioxidant capacity in affected humans and animals was significantly lower than in the controls (*P < 0.01*). The serum protein parameters in affected humans and animals were significantly reduced (*P* < 0.01). It was therefore concluded that heavy metal contamination has caused serious harm to sheep in this area. The heavy metal concentrations in food and grain also pose a significant risk to human health in the Chinese Wumeng mountain area.

## Introduction

The Wumeng mountain area is located in the Yunnan-Guizhou Plateau of China, where the three provinces of Guizhou, Yunnan, and Sichuan meet, and is an important pasture land for sheep [1]. Sheep farming is vital to the production system in the Wumeng mountain area. The animals provide meat, wool, and hides for local people [2]. During the past 10 years, lead, cadmium, copper, and zinc concentrations in air, water, soils, forages, and foods (corn, rice, and wheat) have been increasing in the region. In terms of the potential adverse effects on human and animal health, lead, cadmium, and arsenic have caused most concern [3–4], because they are readily transferred though food-chains and are not known to serve any essential biological function [5]. Industrial emissions of cadmium are the largest source of environmentally hazardous amounts of cadmium [6–7]. The most polluting industries are those associated with mining and smelting, followed by manufacturing, with losses of heavy metals from manufactured products during use and when discarded. The reclamation and use of waste products contaminated with cadmium can also lead to pollution [8–9]. Lead is considered to be a major environmental contaminant and has been more widely reported as a cause of accidental poisoning in humans and livestock than any other substance [10]. Many papers have reviewed specific aspects of heavy metal lead toxicology in humans [10–12] and livestock [7, 13–14]. Lead as an environmental contaminant is often combined with cadmium. Both metals generate similar health effects, and therefore the effects are additive [15–16].

The Wumeng mountain area is also an important production base of non-ferrous metals in China. The area has extensive heavy metal reserves, characterized by large quantities of ores containing zinc, copper, and lead [2]. A large number of industrial enterprises were established for the purpose of lead, zinc, copper, and polymetallic extraction in the 2010s. After a few years of intensive development, metallurgical industries occupied a very wide area of former pasture and farmland [1]. A number of sheep grazing on pastures in the vicinity died after the smelters went into operation. All of the affected sheep were characterized by anemia, emaciation, anorexia, depression, and weakness. The body temperature, respiratory rate, and heart rate of affected animals were normal. In the most severely affected area, 48.36% of sheep were affected and the mortality rate reached 70.67%. Local farmers have suffered seriously from heavy metal contamination, which has caused major economic losses and become a serious scourge. Heavy metals have entered the bodies of local farmers through the food chain, interfering with the normal functions of the body and generating serious harm to their health. However, little research has been undertaken on the movement of heavy metal contaminants in the environment, the effects on animal health, and particularly the effects on local farmers following the passage of heavy metals through the food chain.

The aim of the study was to determine the relationship between the deaths of sheep and the possible environmental impact of the local metallurgical industry, and to investigate the effect of heavy metal contamination on human health in the Wumeng mountain area of China.

## Materials and methods

### Ethics statement

Sheep used in these studies were cared as per outlined in the Guide for the Care and Use of Animals in Agricultural Research and Teaching Consortium. Samples collections in animals were approved by the Institute of Zoology, Chinese Academy of Sciences, Institutional Animal Care and Use Committee (Project A0066).

The human subjects research was approved by the State Engineering Technology Institute for Karst Desertification Control Human Subjects Protection Committee approved our methods, and all participants gave informed consent. All patients provided written informed consent.

### Study area

The area studied was located in the Yunnan-Guizhou Plateau of China (25°49’–28°35’N, 102°45’–105°17’E), with an average elevation of 2200 m above sea level. The annual precipitation is 960 mm and average atmospheric temperature is 10–12°C. The polluted area was located in Hezhang County on the Chinese Yunnan-Guizhou Plateau (26°36’–27°26’N, 103°36’–104°45’E). The control area was located in Dushan County on the Yunnan-Guizhou Plateau (26°29’–27°28’N, 103°33’–104°45’E). The grassland vegetation is mainly Puccinellia (*Chinam poensis ohuji*), Siberian Nitraria (*Nitraria sibirica Pall*), floriated astragalus (*Astragalus floridus*), poly-branched astragals (*Astragalus polycladus*), falcate whin (*Oxytropis falcate*), ewenki automomous banner (*Elymus nutans*), common leymus (*Leymus secalinus*), and june grass (*Koeleria cristata*). Most of the plants are herbaceous and are good resources for grazing animals. The grain crops grown in the are mainly maize, wheat, and rice.

### Selected humans and animals

Fifteen affected sheep aged 2–3 years, were selected from polluted pasture land in Hezhang County in the Wumeng mountain area. All 15 sheep displayed obvious signs of poisoning, including anemia, emaciation, anorexia, and weakness. Fifteen healthy sheep aged 2–3 years, were selected from healthy pasture land in Dushan County in Guizhou province in China. A clinical examination revealed that all animals were in good health.

Forty human volunteers aged 20–30 years, were selected to participate in the study Twenty were selected from a polluted area in Hezhang County in the Wumeng mountain area. The other 20 were selected from an uncontaminated area in Dushan County on the Yunnan-Guizhou Plateau.

### Sample collection

Animal blood samples (15 ml) were obtained from the jugular vein of all sheep. Human blood samples (15 ml) were obtained from the arm vein of ten human volunteers, using 1% sodium heparin as an anticoagulant and stored at −10°C prior to the analysis of heavy metals. Wool was taken from the neck of all sheep. Hair was taken from the head of 20 human volunteers, washed and degreased as described by [17] and kept in a desiccator over silica gel prior to analysis.

All sheep were killed by exsanguination and samples of at least 40 g were taken from the lobus caudatus of the liver, renal cortex of the right kidney, left ventricle of the heart, spleen, lobes of the lung, a gluteal muscle of the left posterior limb, last rib, radius of the left forelimb, and molar teeth. These samples were packed in labelled plastic bags and immediately transported to the laboratory. Visible fat, connective tissue, and major blood vessels were removed from the soft tissues, which were then dried at 80° for 48 h, ground by a mortar, passed through a 0.5 mm sieve, and then stored in a desiccator over silica gel.

Samples of water, soil, and herbage were taken at 48 sampling sites situated at distances of 50–30000 m from the zinc smelter (Fig 1). Multiple small portions of herbaceous vegetation were cut from the pasture in this area and mixed together. To reduce soil contamination, the forage samples were cut at 1 cm above ground level. The herbage samples were dried at 80°C for 48 h and ground by a mortar to facilitate chemical analysis [18–19]. Soil samples were taken from the surface layer (0–30 cm) of the pastures, using a 30-mm diameter cylindrical corer. Soil samples were dried at 80°C for 48 h and passed through a 5 mm sieve. Water samples for irrigating the pasture and farmland from the smelters and food samples (grains of rice, corn, and wheat) were also collected from the farmland of a local farmer. Food samples were dried at 80°C for 48 h, ground by a mortar, passed through a 0.5 mm sieve and then stored in a desiccator over silica gel (Fig 2). Samples of water, soil, forage, and food for use as controls were collected from Dushan County on the Yunnan-Guizhou Plateau.

**Fig 1.**
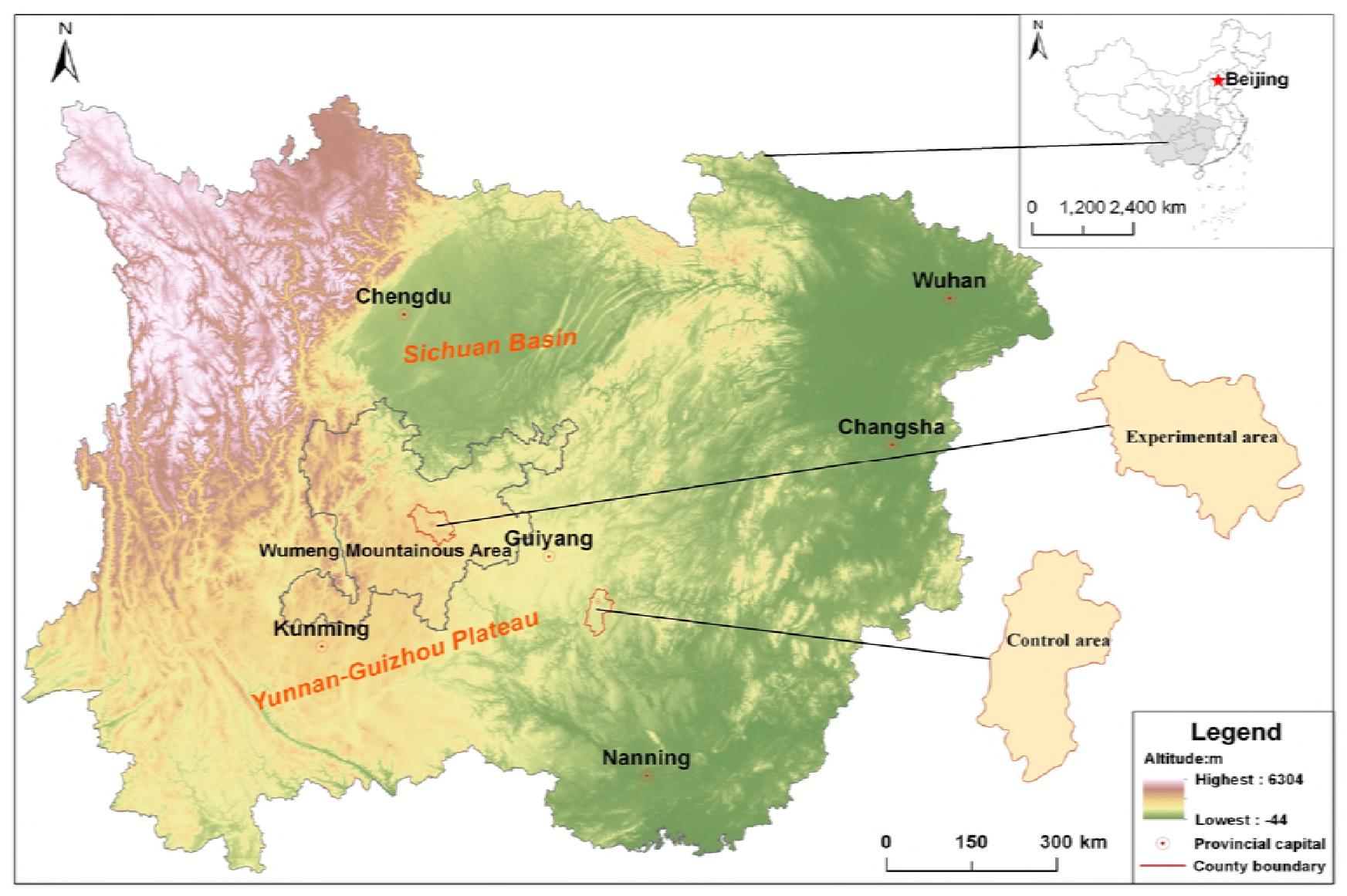
Study area on the Yuannan-Guizhou Plateau.

**Fig 2.**
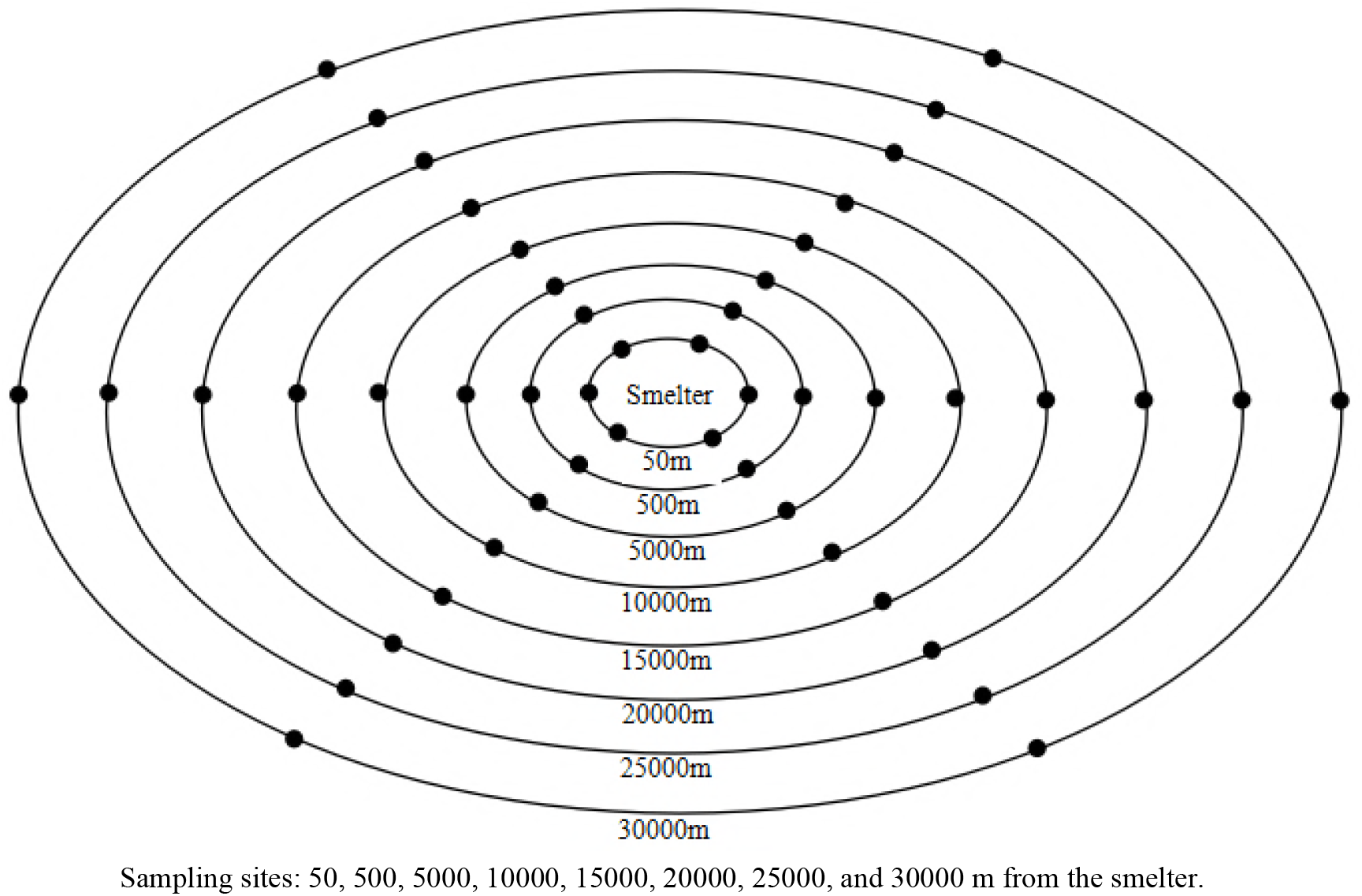
Schematic of the sampling distribution for soil, water, forage, and food samples from the affected pasture.

### Hematological and biochemical examination

Hemoglobin (Hb), packed cell volume (PCV), red blood cell (RBC), white blood cell (WBC), neutrophil, lymphocyte, eosinophil, basophil, and monocyte levels were determined using an automated hematology analyzer (SF-3000, Sysmex-Toa Medical Electronic, Kobe, Japan). Ceruloplasmin (Cp), lactate dehydrogenase (LDH), aspartate aminotransferase (AST), alanine aminotransferase (ALT), alkaline phosphatase (AKP), creatinine CRT), cholesterol (Chol), blood urea nitrogen (BUN), glutathione peroxidase (GSH-Px), superoxide dismutase (SOD), Malondialdehyde (MDA), total antioxidant capacity (T-AOC), sodium (Na), potassium (K), magnesium (Mg), calcium (Ca), and inorganic phosphorus (IP) levels in serum were determined using an automated biochemical analyzer (Olympus AU 640, Olympus Optical Co., Tokyo, Japan). Serum protein (total protein, albumin, and globulin) electrophoretic studies were performed on cellulose acetate using the EA-4 electrophoresis apparatus (Shanghai Medical Apparatus and Instruments Factory, Shanghai, China). All the serum biochemical values were measured at 25°C.

### Analysis of heavy metals

For each analysis, 1 g of sample (1 ml of blood) was added to 1 ml of hydrogen peroxide and 3 ml of nitric acid. These samples were digested at 180°C in fluoro-plastic vessels in sealed containers, which avoided loss of vapor, and therefore there was no reduction in the volume of the resulting digest. Molybdenum (Mo) was determined using flameless atomic absorption spectrophotometry (3030 graphite furnace with a Zeeman background correction, Perkin-Elmer, Waltham, MA, USA) [18–19]. Lead, cadmium, copper, zinc, and manganese concentrations were determined by atomic absorption spectrophotometry (AA-640, Shimadzu Co., Ltd, Tokyo, Japan). The accuracy of the analytical values was confirmed by reference to certified elemental concentrations in reference materials from the International Atomic Energy Agency certified and National Institute of Standards (Bovine liver SRM 1577a). The limited accuracy of samples with very low concentrations resulted in concentrations below a particular threshold being recorded as ‘trace’, given that zero measurements were difficult to demonstrate.

### Statistical analysis

Data were analyzed using the statistical package for the social sciences (SPSS, version 20.0, Inc., Chicago, IL, USA), and presented in the form of mean ± standard error (SE). Significant differences between groups were assessed using a student’s *t* test, with least significant differences of 1% (*P < 0.01*) or 5% (*P < 0.05*).

## Results

It was found that heavy metal concentrations clearly decreased with increasing distance from the zinc smelter (Fig 3–5). The heavy metal concentrations (lead, cadmium, copper, and zinc) in irrigation water, soils, forages, and foods (corn, rice, and wheat) in affected pastures were markedly higher than those in the control area (*P < 0.01*; Table 1–2). The mean cadmium and lead concentrations in affected pastures exceeded the control levels by 68.71 and 15.66 times in soil, respectively, and 556.15 and 16.61 times in herbage, respectively. Taking into account that these sheep were grazing exclusively in affected pastures, the cadmium and lead ingestion rates were estimated (Table 3). The estimation was based on an average herbage ingestion of 76.3 g (d.w)/kg body wt./day and in the sheep [20]. The ingestion rates were in the range of 3.27–38.47 mg/kg body wt./day and 0.19–3.36 mg/kg body wt./day for lead and cadmium in sheep, respectively.

**Fig 3.**
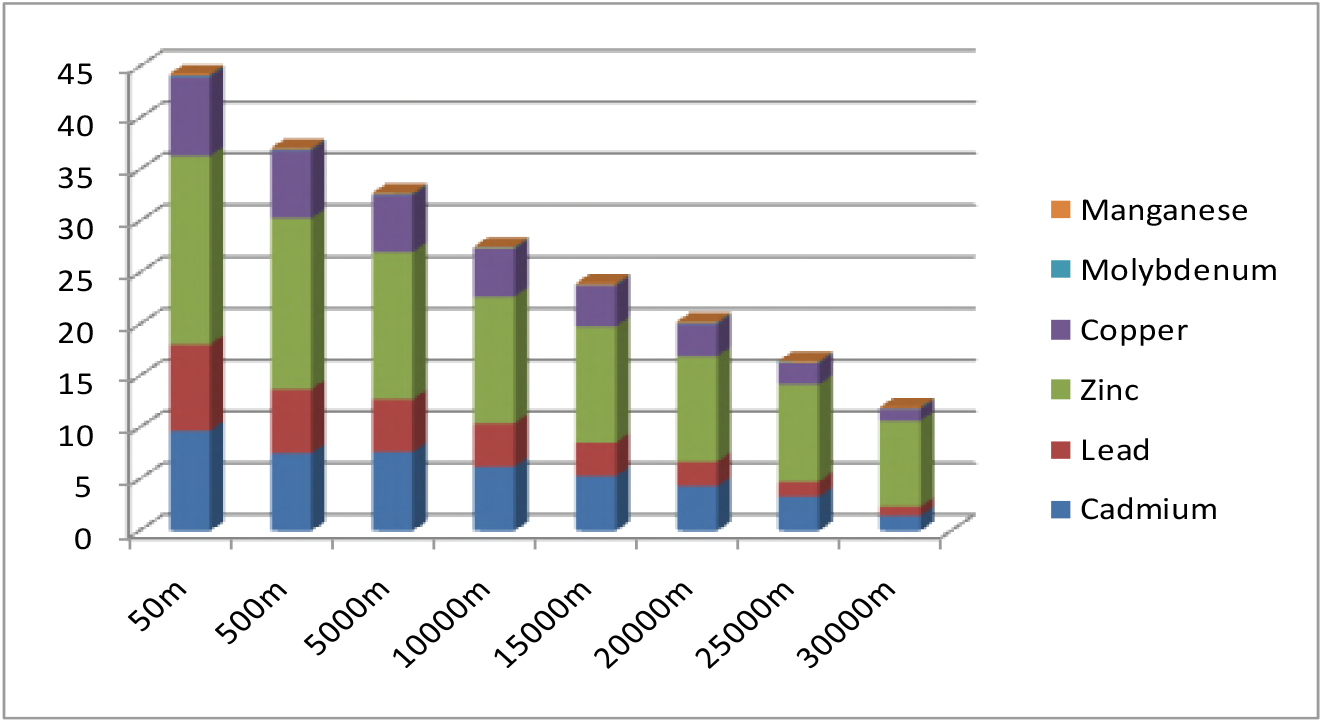
Relationship between heavy metal concentrations and the distance of sampling sites from the smelter in irrigation water (mg/L)

**Fig 4.**
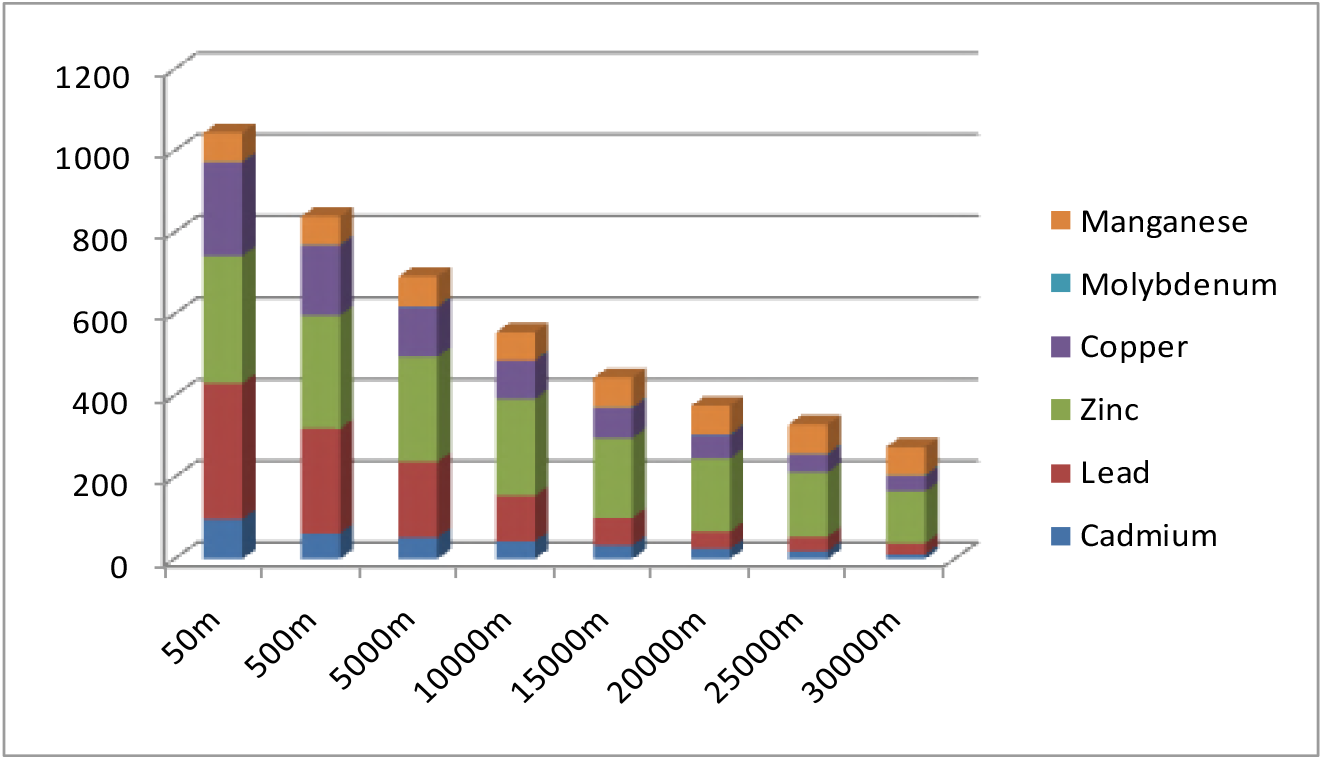
Relationship between heavy metal concentrations and the distance of sampling sites from the smelter in soils (mg/kg DM)

**Fig 5.**
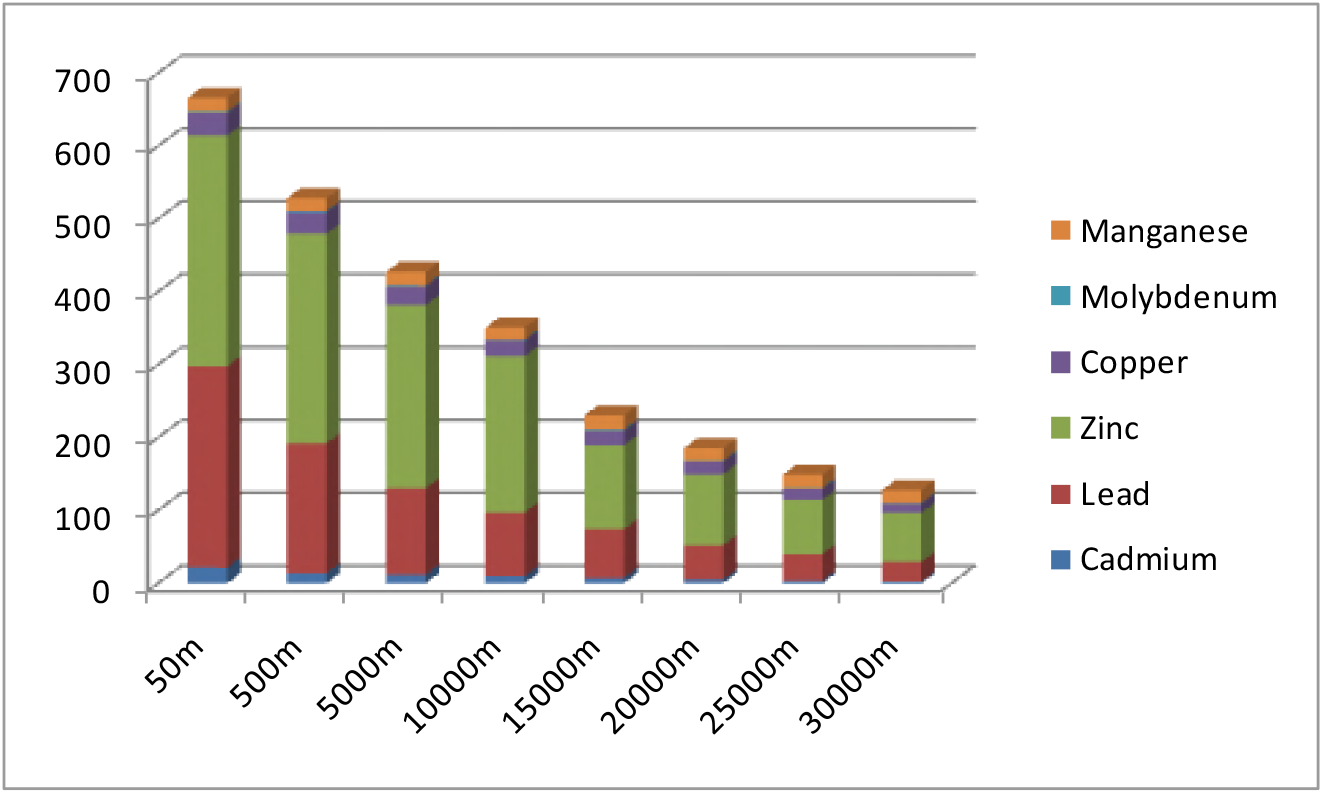
Relationship between heavy metal concentrations and the distance of sampling sites from the smelter in forages (mg/kg DM)

**Table 1.**
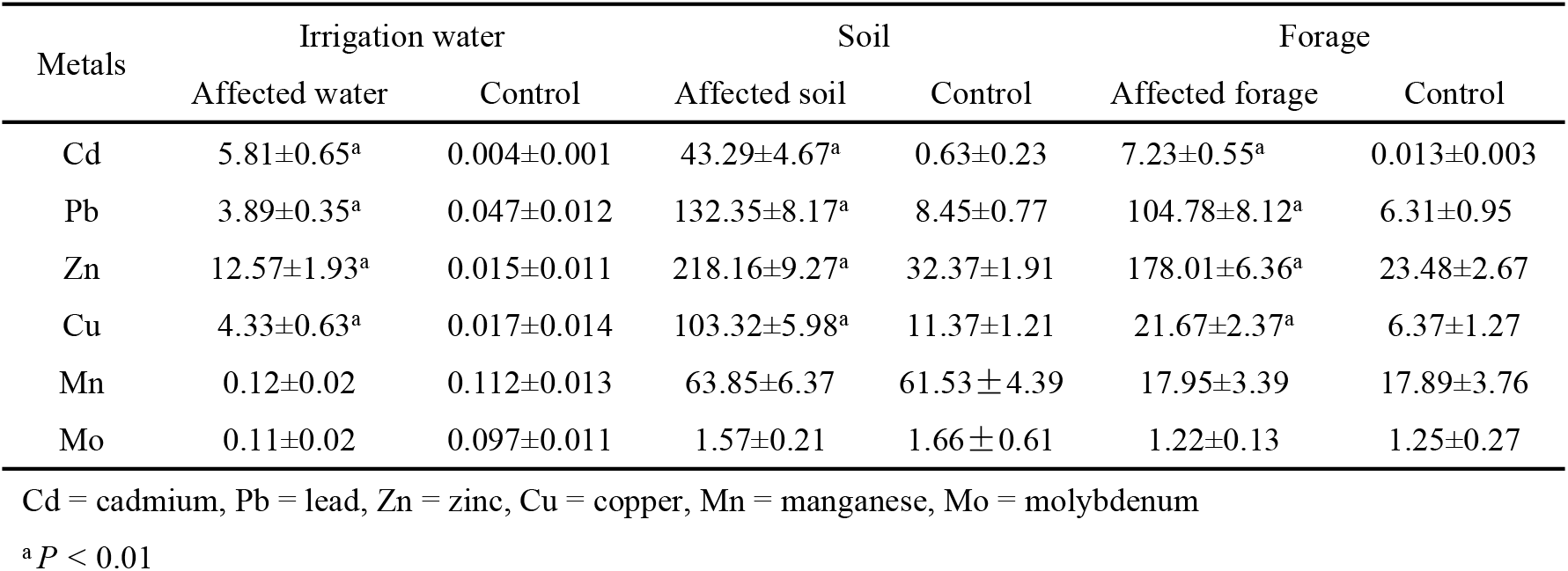
Metal concentrations in soil, irrigation water, and herbage in all samples (mg/kg)

**Table 2.**
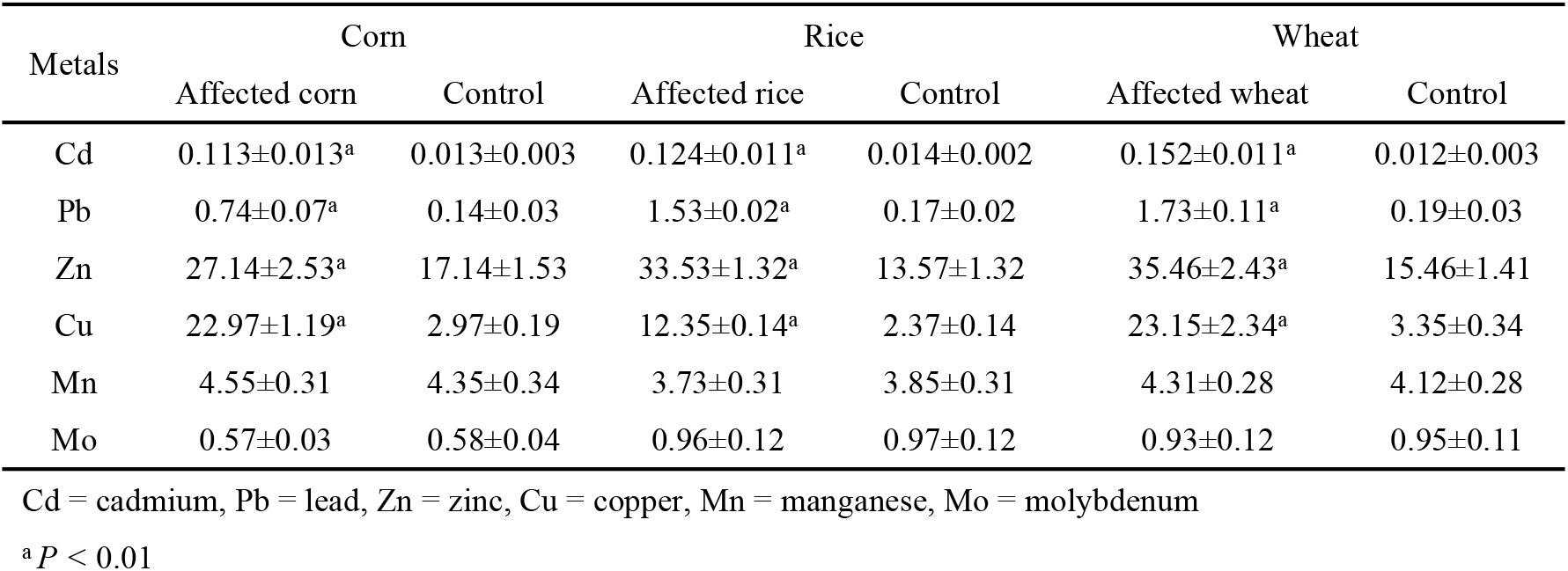
Metal concentrations in corn, rice, and wheat in all samples (mg/kg)

| Metals | Corn |  | Rice |  | Wheat |  |
| --- | --- | --- | --- | --- | --- | --- |
|  | Affected corn | Control | Affected rice | Control | Affected wheat | Control |
| Cd | 0.113±0.013<sup>a</sup> | 0.013±0.003 | 0.124±0.011<sup>a</sup> | 0.014±0.002 | 0.152±0.011<sup>a</sup> | 0.012±0.003 |
| Pb | 0.74±0.07<sup>a</sup> | 0.14±0.03 | 1.53±0.02<sup>a</sup> | 0.17±0.02 | 1.73±0.11<sup>a</sup> | 0.19±0.03 |
| Zn | 27.14±2.53<sup>a</sup> | 17.14±1.53 | 33.53±1.32<sup>a</sup> | 13.57±1.32 | 35.46±2.43<sup>a</sup> | 15.46±1.41 |
| Cu | 22.97±1.19<sup>a</sup> | 2.97±0.19 | 12.35±0.14<sup>a</sup> | 2.37±0.14 | 23.15±2.34<sup>a</sup> | 3.35±0.34 |
| Mn | 4.55±0.31 | 4.35±0.34 | 3.73±0.31 | 3.85±0.31 | 4.31±0.28 | 4.12±0.28 |
| Mo | 0.57±0.03 | 0.58±0.04 | 0.96±0.12 | 0.97±0.12 | 0.93±0.12 | 0.95±0.11 |

Cd = cadmium, Pb = lead, Zn = zinc, Cu = copper, Mn = manganese, Mo = molybdenum
<sup>a</sup> $P < 0.01$

**Table 3.**
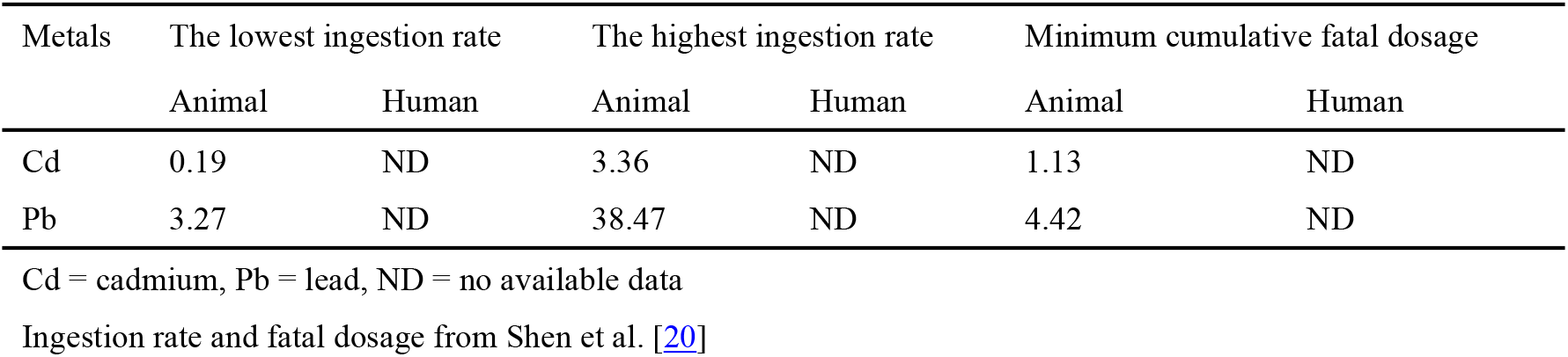
The ingestion rates and fatal dosage in animals and humans (mg/kg body wt./day)

The lead, cadmium, zinc, and copper concentrations in wool, blood, heart, lung, liver, muscle, spleen, and bone of affected sheep were markedly higher than those in healthy animals (*P < 0.01*) (Table 4–5). Lead and cadmium concentrations were mainly accumulated in the kidney, liver, and skeleton of affected sheep in this area. Lead, cadmium, zinc, and copper concentrations in blood and hair samples from affected local farmers were significantly higher than those in the controls (*P < 0.01*; Table 6).

**Table 4.** Metal concentrations in wool and blood samples from sheep (mg/kg)

| Metals | Wool |  | Blood |  |
| --- | --- | --- | --- | --- |
|  | Affected animal | Control | Affected animal | Control |
| Cd | 2.28±0.13<sup>a</sup> | 0.36±0.03 | 0.391±0.02<sup>a</sup> | 0.013±0.01 |
| Pb | 3.76±0.21<sup>a</sup> | 1.16±0.12 | 0.411±0.021<sup>a</sup> | 0.042±0.03 |
| Zn | 119.97±7.23<sup>a</sup> | 87.61±5.97 | 14.82±0.97<sup>a</sup> | 9.81±2.16 |
| Cu | 9.87±0.23<sup>a</sup> | 3.73±0.57 | 1.74±0.07<sup>a</sup> | 0.79±0.12 |
| Mn | 4.89±0.47 | 4.63±0.36 | 0.38±0.03 | 0.35±0.02 |
| Mo | 2.17±0.12 | 2.18±0.13 | 0.19±0.07 | 0.15±0.05 |

Cd = cadmium, Pb = lead, Zn = zinc, Cu = copper, Mn = manganese, Mo = molybdenum
<sup>a</sup> $P < 0.01$

**Table 5.** Metal concentrations in heart, lung, liver, kidney, muscle, spleen, rib, radius, and tooth samples from sheep

| Metals | Heart |  | Lung |  | Liver |  |
| --- | --- | --- | --- | --- | --- | --- |
|  | Affected animal | Control | Affected animal | Control | Affected animal | Control |
| Cd | 0.39±0.13<sup>a</sup> | 0.21±0.12 | 3.12±0.25<sup>a</sup> | 0.67±0.13 | 7.62±0.56<sup>a</sup> | 0.47±0.02 |
| Pb | 4.76±0.67<sup>a</sup> | 1.34±0.23 | 4.45±0.38<sup>a</sup> | 1.55±0.27 | 16.37±1.57<sup>a</sup> | 0.73±0.05 |
| Zn | 137.63±6.56<sup>a</sup> | 103.92±7.23 | 143.87±8.67<sup>a</sup> | 107.82±7.63 | 233.93±8.72<sup>a</sup> | 137.79±8.57 |
| Cu | 29.81±3.43<sup>a</sup> | 14.72±0.93 | 25.23 ±2.32<sup>a</sup> | 11.67±1.69 | 258.17±7.93<sup>a</sup> | 117.67±7.73 |
| Mn | 3.67±0.53 | 3.52±0.23 | 3.82±0.27 | 3.83±0.31 | 4.73±0.72 | 4.56±0.59 |
| Mo | 1.32±0.11 | 1.33±0.17 | 1.53±2.46 | 1.68±1.67 | 1.43±0.13 | 1.39±0.18 |
| Metals | Kidney |  | Muscle |  | Spleen |  |
|  | Affected animal | Control | Affected animal | Control | Affected animal | Control |
| Cd | 27.37±1.53<sup>a</sup> | 1.73±0.29 | 0.82±0.03<sup>a</sup> | 0.18±0.04 | 0.57±1.71<sup>a</sup> | 0.33±0.05 |
| Pb | 38.57±2.87<sup>a</sup> | 0.97±0.13 | 2.85±0.52<sup>a</sup> | 0.87±0.03 | 3.87±8.53<sup>a</sup> | 0.93±0.11 |
| Zn | 157.16±7.43<sup>a</sup> | 113.75±7.69 | 141.13±7.62<sup>a</sup> | 113.56±9.87 | 137.93±9.17<sup>a</sup> | 118.62±9.86 |
| Cu | 27.21±3.27<sup>a</sup> | 11.32±1.73 | 8.97±0.97<sup>a</sup> | 5.83±0.96 | 9.52±0.57<sup>a</sup> | 4.47±0.56 |
| Mn | 4.68±0.57 | 4.73±0.85 | 3.83±0.57 | 3.82±0.67 | 2.56±0.37 | 2.61±0.29 |
| Mo | 1.87±0.42 | 1.79±0.92 | 1.28 ±0.48 | 1.36±0.68 | 0.79±0.14 | 0.37±0.03 |
| Metals | Rib |  | Radius |  | Tooth |  |
|  | Affected animal | Control | Affected animal | Control | Affected animal | Control |
| Cd | 4.77±0.23<sup>a</sup> | 1.33±0.27 | 5.62±0.73<sup>a</sup> | 1.78±0.34 | 5.57±0.54<sup>a</sup> | 1.43±0.23 |
| Pb | 14.53±2.73<sup>a</sup> | 4.37±0.23 | 22.53±2.51<sup>a</sup> | 6.57±0.73 | 27.82±2.73<sup>a</sup> | 5.11±0.95 |
| Zn | 97.83±3.43<sup>a</sup> | 77.75±3.69 | 117.13±5.62<sup>a</sup> | 83.56±9.87 | 137.37±7.59<sup>a</sup> | 113.97±7.61 |
| Cu | 8.21±0.57<sup>a</sup> | 4.32±0.73 | 11.17±0.77<sup>a</sup> | 4.83±0.96 | 8.53±1.27<sup>a</sup> | 5.73±0.67 |
| Mn | 2.17±0.32 | 2.09±0.32 | 1.37 ±0.48 | 1.36±0.68 | 1.76±0.31 | 1.69±0.27 |
| Mo | 3.68±0.35 | 3.73±0.35 | 4.23±0.53 | 3.82±0.67 | 4.22±0.53 | 4.17±0.76 |

Cd = cadmium, Pb = lead, Zn = zinc, Cu = copper, Mn = manganese, Mo = molybdenum

**Table 6.** Metal concentrations in hair and blood samples from humans (mg/kg)

| Metals | Hair |  | Blood |  |
| --- | --- | --- | --- | --- |
|  | Affected human | Control | Affected human | Control |
| Cd | 1.88±0.12<sup>a</sup> | 0.37±0.02 | 0.29±0.01<sup>a</sup> | 0.013±0.01 |
| Pb | 2.71±0.33<sup>a</sup> | 1.23±0.31 | 0.31±0.03<sup>a</sup> | 0.042±0.03 |
| Zn | 99.63±7.71<sup>a</sup> | 67.63±6.76 | 14.82±0.81<sup>a</sup> | 6.81±2.16 |
| Cu | 8.82±0.27<sup>a</sup> | 3.21±0.53 | 1.34±0.007<sup>a</sup> | 0.69±0.01 |
| Mn | 4.17±0.37 | 4.27±0.32 | 0.37±0.12 | 0.35±0.12 |
| Mo | 2.23±0.15 | 2.19±0.11 | 0.17±0.03 | 0.15±0.05 |

Cd = cadmium, Pb = lead, Zn = zinc, Cu = copper, Mn = manganese, Mo = molybdenum
<sup>a</sup> $P < 0.01$

Hematological parameters in affected farmers and sheep are given in Table 7. Compared with healthy farmers and sheep, hemoglobin levels and packed cell volume were markedly reduced (*P < 0.01*). These abnormal blood indices indicated a hypochromic microcytic anemia in affected humans and animals. Serum biochemical parameters in affected humans and animals are given in Table 8. Compared with healthy humans and animals, creatinine, lactate dehydrogenase, superoxide dismutase, and glutathione peroxidase activities were significantly reduced (*P < 0.01*). Serum total antioxidant capacity levels in affected humans and animals were significantly lower than those in the controls (*P < 0.01*). Malondialdehyde levels in serum from affected humans and animals were significantly higher than those in the controls. Serum protein parameters in affected humans and animals are given in Table 9. Compared with healthy humans and animals, the total protein, albumin, α-globulin, β-globulin, and γ-globulin levels were significantly reduced (*P < 0.01*).

**Table 7.** Hematological parameters in animals and humans

| Parameters | Animal |  | Human |  |
| --- | --- | --- | --- | --- |
|  | Affected animal | Control | Affected human | Control |
| Hb (g/L) | 97.83±6.56<sup>a</sup> | 124.56±9.65 | 87.65±6.56<sup>a</sup> | 131.78±9.57 |
| RBC ($10^{12}/L$) | 12.36±2.65 | 12.15±3.66 | 4.31±0.56 | 4.53±0.43 |
| PCV (%) | 31.21±4.23<sup>a</sup> | 39.13±3.15 | 42.57±4.36<sup>a</sup> | 45.17±6.56 |
| WBC($10^9/L$) | 6.38±1.97 | 6.56±2.13 | 5.18±0.77 | 5.36±0.63 |
| Neutrophils (%) | 56.98±9.87 | 57.56±9.18 | 63.98±6.81 | 65.46±5.38 |
| Lymphocytes (%) | 29.79±6.31 | 30.17±6.19 | 35.72±3.31 | 33.27±3.17 |
| Eosinophils (%) | 6.42±0.58 | 6.56±0.37 | 7.12±0.98 | 7.46±0.87 |
| Basophils (%) | 0.48±0.23 | 0.51±0.32 | 0.52±0.27 | 0.57±0.33 |
| Monocytes (%) | 7.13±0.64 | 6.97±0.67 | 5.76±0.57 | 5.36±0.43 |

Hb = hemoglobin, PCV = packed cell volume, RBC = red blood cells, WBC = white blood cells
<sup>a</sup> $P < 0.01$

**Table 8.** Serum biochemical parameters in animals and humans

| Parameters | Animal |  | Human |  |
| --- | --- | --- | --- | --- |
|  | Affected animal | Control | Affected human | Control |
| Cp (mg/L) | 51.37±11.67 | 51.76±11.26 | 56.36±6.29 | 55.37±6.17 |
| LDH (U/L) | 2.51±0.32<sup>a</sup> | 3.67±0.43 | 2.73±0.37<sup>a</sup> | 3.57±0.23 |
| AKP (U/L) | 76.78±13.98 | 77.89±16.56 | 87.18±7.96 | 86.78±7.23 |
| AST (U/L) | 38.73±3.36 | 37.93±4.57 | 38.27±4.22 | 37.73±3.31 |
| ALT (U/L) | 13.36±3.27 | 13.88±3.77 | 22.31±3.18 | 22.36±3.36 |
| BUN (U/L) | 6.62±1.37 | 6.57±1.76 | 6.57±0.97 | 6.33±0.61 |
| SOD (U/L) | 111.35±8.73<sup>a</sup> | 157.67±9.69 | 123.23±9.77<sup>a</sup> | 143.37±9.38 |
| GSH-Px (U/L) | 287.67±9.38<sup>a</sup> | 389.56±9.77 | 233.62±7.18<sup>a</sup> | 353.72±7.49 |
| CAT (U/L) | 16.63±1.25 | 16.92±1.63 | 19.68±1.73 | 19.63±1.52 |
| T-AOC (U/L) | 3.01±0.13<sup>a</sup> | 4.32±0.17 | 3.31±0.14<sup>a</sup> | 4.37±0.12 |
| CRT (U/L) | 257.87±13.27<sup>a</sup> | 318.67±7.83 | 277.51±13.89<sup>a</sup> | 336.52±11.35 |
| Chol (U/L) | 2.77±0.31 | 2.81±0.36 | 2.83±0.37 | 2.79±0.31 |
| MDA (nmol/L) | 11.37±0.38<sup>a</sup> | 6.36±0.15 | 12.53±0.39<sup>a</sup> | 7.23±0.57 |
| K (mmol/L) | 3.99±0.37 | 4.03±0.58 | 3.75±0.38 | 3.35±0.37 |
| Na (mmol/L) | 129.35±5.89 | 121.76±8.37 | 116.35±5.33 | 113.37±4.31 |
| Ca (mmol/L) | 2.28±0.21 | 2.31±0.23 | 2.52±0.17 | 2.72±0.19 |
| IP (mmol/L) | 1.89±0.32 | 1.85±0.26 | 1.63±0.12 | 1.57±0.17 |
| Mg (mmol/L) | 0.89±0.11 | 0.91±0.21 | 0.73±0.03 | 0.81±0.04 |

Cp = ceruloplasmin, LDH = lactate dehydrogenase, AKP = alkaline phosphatase, AST = aspartate aminotransferase, ALT = alanine aminotransferase, BUN = blood urea nitrogen, SOD = superoxide dismutase, GSH-Px = glutathione peroxidase, CAT = catalase, MDA = malondialdehyde, T-AOC = total antioxidant capacity, CRT = creatinine, Chol = cholesterol, K = potassium, Na = sodium, Ca = calcium, IP = inorganic phosphorus, Mg = magnesium.
<sup>a</sup> $P < 0.01$

**Table 9.**
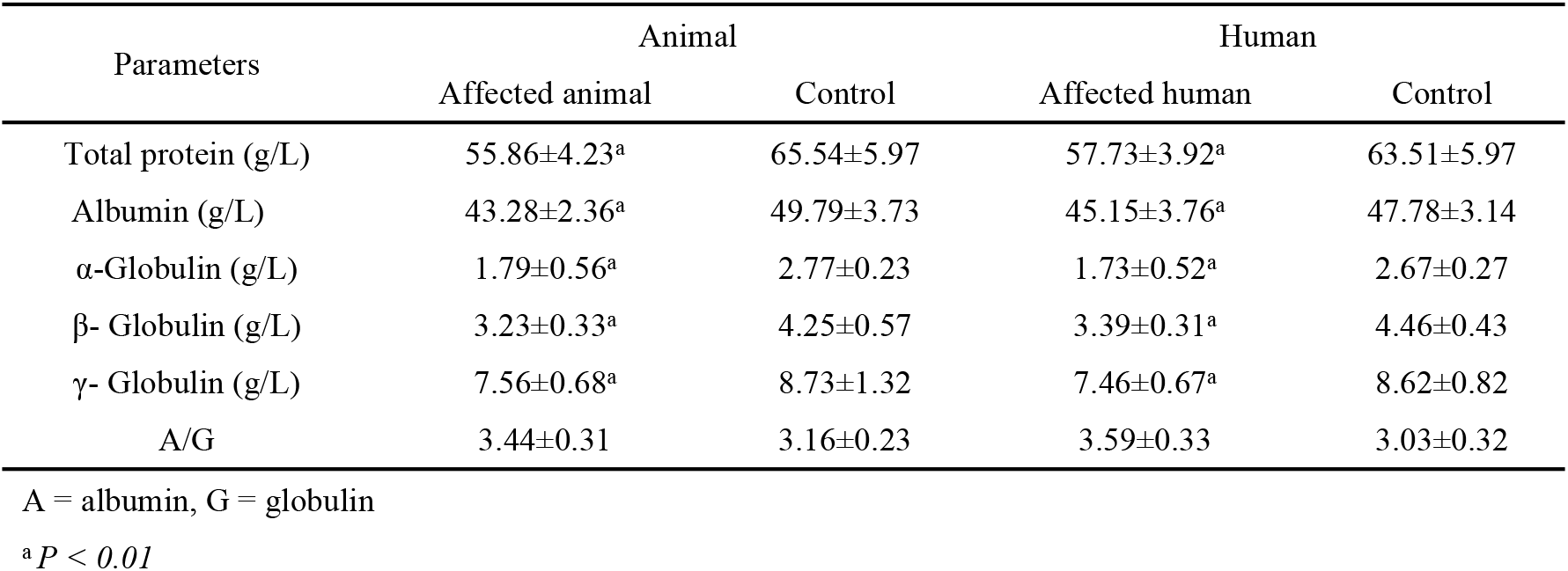
Serum protein concentrations in animals and humans

| Parameters | Animal |  | Human |  |
| --- | --- | --- | --- | --- |
|  | Affected animal | Control | Affected human | Control |
| Total protein (g/L) | 55.86±4.23<sup>a</sup> | 65.54±5.97 | 57.73±3.92<sup>a</sup> | 63.51±5.97 |
| Albumin (g/L) | 43.28±2.36<sup>a</sup> | 49.79±3.73 | 45.15±3.76<sup>a</sup> | 47.78±3.14 |
| α-Globulin (g/L) | 1.79±0.56<sup>a</sup> | 2.77±0.23 | 1.73±0.52<sup>a</sup> | 2.67±0.27 |
| β- Globulin (g/L) | 3.23±0.33<sup>a</sup> | 4.25±0.57 | 3.39±0.31<sup>a</sup> | 4.46±0.43 |
| γ- Globulin (g/L) | 7.56±0.68<sup>a</sup> | 8.73±1.32 | 7.46±0.67<sup>a</sup> | 8.62±0.82 |
| A/G | 3.44±0.31 | 3.16±0.23 | 3.59±0.33 | 3.03±0.32 |

A = albumin, G = globulin
<sup>a</sup> $P < 0.01$

## Discussion

The contamination of pasture and farmland with heavy metals (cadmium, lead, copper, and zinc) may occur in the vicinity of smelters. The maximum tolerable dietary content of cadmium and lead has been set at 0.5 and 30 mg/kg for livestock, respectively [13]. Cadmium and lead concentrations in soil, herbage, and irrigated water samples in affected pastures significantly exceeded these levels (*P < 0.01*). The lead, cadmium, copper, and zinc concentrations in these samples decreased markedly with increasing distance from the zinc smelter in the Wumeng mountain area. This tendency has been observed in other studies of heavy metal contamination in areas surrounding smelting industries [21] and indicates the aeolian dispersion of particles containing heavy metals (lead, cadmium, copper, and zinc) from these smelters are a source of heavy metal contamination in soils [22–23]. Waste water discharged from the zinc smelter has been used to irrigate surrounding pastures and farmland in the Wumeng mountain area and was therefore also a source of heavy metals (cadmium, lead, copper, and zinc) in the agricultural soils of the study area.

Lead, as an environmental contaminant, is often associated with cadmium, with both elements having similar properties and their health effects being additive [21, 24]. In this study, lead, cadmium, copper, and zinc concentrations in the tissues (wool, blood, heart, lung, liver, muscle, spleen, and bone) of affected sheep were markedly higher than those in the controls (*P < 0.01*). Hypochromic microcytic anemia was evident in affected animals. It is generally believed that high skeletal lead and cadmium concentrations are a characteristic of chronic exposure to lead and cadmium [25–27]. The cortex of the kidney in the affected sheep contained higher lead and cadmium concentrations than the liver. Cadmium and lead are also accumulated in the bones of affected sheep. These results are consistent with previous studies of livestock that indicate that these tissues are the critical organs for lead and cadmium accumulation [28–30]. It was therefore concluded that heavy metal contamination due to industrial activities in the Wumeng mountain area had resulted in serious harm to sheep health.

As a result of activities conducted in the zinc smelters in the Wumeng mountain area, a large increase in cadmium, lead, copper, and zinc concentrations was observed in the surrounding soils and herbage. Taking into account that the sheep were fed exclusively with forage from this pasture, the ingested heavy metal rates were estimated to be in the range of 3.27–38.47 and 0.19–3.36 mg/kg body wt./day for lead and cadmium, respectively. Registered values for the minimum cumulative fatal dosage for sheep are estimated at 4.42 and 1.13 mg/ body wt./day for lead and cadmium, respectively [14, 31]. Therefore, the ingestion of forages growing in this pasture, especially in the sites closest to the zinc smelters constitutes a clear cadmium and lead toxicity hazard for livestock. Lead and cadmium intake levels in the affected pasture surpassed the fatal dosage (*P < 0.01*). As a consequence of the cadmium and lead uptake, the lead and cadmium concentrations in tissues (wool, blood, heart, lung, liver, muscle, spleen, and bone) in affected sheep surpassed the critical and control concentrations. This confirms the potential toxicity of the pasture [32–33]. The sheep were predominantly fed on locally grown fodder or grazed on pasture in the vicinity of zinc smelters and are the primary livestock species exposed to heavy metal contamination in this area. Therefore, a determination of the cadmium, lead, copper, and zinc concentrations in domestic animals in this area is important for assessing the potential effects of pollutants on livestock and for quantifying contaminant uptake by humans.

Lead, cadmium, copper, and zinc concentrations in blood and hair samples from the local farmers in the affected area were also significantly higher than those in the controls (*P < 0.01*). Hypochromic microcytic anemia was also evident in affected humans. The levels of superoxide dismutase, glutathione peroxidase, and total antioxidant capacity are given in Table 8. Serum total antioxidant capacity is an integrative index used to reflect the antioxidant capacity of the body [34–36]. Little is known about the effects of lead and cadmium on the total antioxidant capacity of sheep. Our results indicate that the total antioxidant capacity levels of affected humans and animals were significantly reduced (*P < 0.01*), and the enhanced peroxidation of lipids in intracellular and extracellular membranes resulted in damage to cells, tissues, and organs. Superoxide dismutase and glutathione peroxidase are important antioxidant enzymes that protect against this process [37]. Superoxide dismutase catalyzes the destruction of the superoxide radical, with potential toxicity arising from dismutation and hydrogen peroxide formation, while glutathione peroxidase catalyzes the conversion of hydrogen peroxide to water and directly reduces tissue injury from lipoperoxidation [34, 38]. A significant decrease in activity of either would therefore induce an increase in free radicals; thus, injuring the corresponding tissues. The results show that the superoxide dismutase and glutathione peroxidase activities in the serum of affected humans and animals markedly decreased (*P < 0.01*). Thus, it can be seen that heavy metal contamination not only resulted in serious harm to sheep health, but also entered the human body through the food chain and interfered with the normal functions of the body, resulting in serious harm to human health.

In this study, the copper and zinc concentrations in soil, herbage, and food were markedly higher in samples from the affected area than those from the control area (*P < 0.01*). In general, the maximum tolerable concentrations in sheep were 25 and 300 mg/kg, for copper and zinc, respectively [13, 39–40]. Thus, it appears that the heavy metal poisoning of the sheep in the pasture was not related to copper and zinc. Whether the copper and zinc concentrations in the soil and herbage affected the absorption of cadmium and lead in animals and humans requires further investigation.

## Acknowledgement

This work was supported by the national key research and development program of China in the 13th five-year plan (2016YFC0502607) and the national natural science foundation of China (41671041).

## Author Contributions

**Conceptualization**: Xiaoyun Shen.

**Data curation:** Kangning Xiong.

**Formal analysis:** Yongkuan Chi.

**Investigation:** Xiaoyun Shen.

**Methodology:** Yongkuan Chi.

**Software:** Yongkuan Chi.

**Supervision:** Kangning Xiong.

**Validation:** Xiaoyun Shen, Yongkuan Chi.

**Visualization:** Yongkuan Chi, Kangning Xiong.

**Writing ± original draft:** Xiaoyun Shen.

